# Proton stress adaptation in acidophilic sulfate-reducing bacteria: insights from *Acididesulfobacillus acetoxydans* for acid mine drainage bioremediation

**DOI:** 10.1101/2025.11.06.686915

**Authors:** Reinier A. Egas, Nicole J. Bale, Michel Koenen, Laura Villanueva, Diana Z. Sousa, Cornelia U. Welte, Irene Sánchez-Andrea

**Author notes:** Address correspondence to Reinier A. Egas.

## Abstract

Acid mine drainage (AMD) waters are a global environmental threat due to their extremely low pH (<3) and high metal loads. Acidophilic sulfate-reducing bacteria (aSRB) can mitigate AMD by reducing sulfate to sulfide, a proton-consuming process that also precipitates metals as metal sulfides. Although sulfate reduction has been observed in AMD waters, most characterized aSRB are only moderately acidophilic and originate from protected microniches. Here, we examined the pH tolerance and proton stress adaptation of the complete organic acid-oxidizing aSRB *Acididesulfobacillus acetoxydans*. Continuous chemostat cultivations were operated across a pH gradient, achieving steady states from pH 5.0 (optimum) down to 2.9, with microcosms showing metabolic activity even at pH 2.5 which is typical AMD-acidity. Transcriptomic profiles remained remarkably stable across conditions, except for upregulation of the K^+^-transporting ATPase (*kdpABC*) at lower pH, suggesting an increased reliance on the chemiosmotic gradient to impede proton influx. Lipid analysis revealed increased core lipid saturation, mid-chain methylation and a shift in priming precursors from leucine to valine at low pH, indicating reduced membrane permeability and more energy-efficient biosynthetic pathways. These adaptations impede proton entry demonstrating adaptation of aSRB to AMD-like acidity and removes the critical pH bottleneck for AMD bioremediation and metal recovery.

**Synopsis:** This study shows acidophilic sulfate-reducing bacteria can adapt to AMD-like acidity while retaining metabolic activity, underscoring their potential for AMD bioremediation and biotechnology.

## Introduction

Acid mine and acid rock drainage (AMD) are causing globally significant environmental problems characterized by extremely low pH (<3) and high concentrations of sulfate and toxic metals such as copper, nickel and zinc ^1^. If untreated, AMD discharges cause severe ecological damage and contaminate a vast amount of downstream water bodies ^2–4^. Effective remediation requires in general three outcomes: pH neutralization, sulfate reduction, and metal removal ^5,6^.

Bioremediation is the active biological mitigation process to treat AMD: several biological processes generate alkalinity and can contribute to bioremediation, including oxygenic photosynthesis, methanogenesis, denitrification, ammonification, iron, sulfur and sulfate reduction ^1,7^. Among these, sulfate reduction is particularly promising as it not only consumes protons but also produces hydrogen sulfide, which precipitates metals as sparingly soluble metal sulfides ^8,9^. Industrial processes employing neutrophilic sulfate-reducing bacteria (SRB) have operated since the 1990s ^10^, but their application to AMD is limited mainly due to low pH and high metal loads, necessitating costly pre-treatment (*e.g.*, lime addition) or process separation (*e.g.*, separate contactor tank) ^5,11^.

Novel applications of acidophilic and acidotolerant SRB (aSRB) circumvent these disadvantages by handling AMD without pre-treatment. In addition to bioremediation, this has the additional benefit of allowing selective recovery of metals ^12–16^. Moderate acidophiles grow optimally near pH 5.0, while extreme acidophiles can thrive at pH ≤ 3 ^17^. Although (a)SRB have been detected in AMD water columns, characterized aSRB originate from sediments or protected microniches where these are shielded from pH differences and high metal concentrations ^16–18^ (Table S1). Curiously, reported *in situ* activity of sulfate reduction occurs at lower pH than the growth thresholds of cultured isolates ^18–20^. This discrepancy indicates that the mechanisms and tolerance of proton stress adaptation in aSRB remain poorly understood.

Proton stress resistance results from an interplay between two defensive lines: (i) limiting or impeding proton influx across the membrane and (ii) exporting or consuming protons that enter the cytoplasm ^17,21,22^. However, the relative contribution of these mechanisms and the effect on sulfate-reduction rates in aSRB has not been resolved. Here, we used continuous chemostat cultures of the complete organic acid oxidizing aSRB *Acididesulfobacillus acetoxydans* ^23,24^ to investigate physiological, transcriptomic, and lipidomic responses to decreasing pH. We demonstrate continuous cultivations and growth at pH 2.9 with activity in subsequent microcosm cultivations at pH 2.5, thereby closing the gap between environmental biosulfidogenesis and laboratory cultivation. The sulfate reduction rates indicate a robust process and high adaptability of aSRB to decreasing pH while the transcriptome and lipid profiles at different pH provide physiological insights into how aSRB tolerate extreme proton stress.

## Materials and Methods

### Strain and media

Cultivations were performed with *A. acetoxydans* DSM 29876^T^. Unless otherwise specified chemicals were obtained from Sigma-Aldrich (Merck KGaA, Darmstadt, Germany). For all cultivations medium was prepared as described previously ^24^ and consisted of (g L^-1^): Na_2_HPO_4_ 2H_2_O, 0.53; KH_2_PO_4_, 0.41; NH_4_Cl, 0.3; NaCl, 0.3; MgCl_2_ · 6H_2_O, 0.1; CaCl_2_ · H_2_O, 0.11 and 0.5 mg L^-1^ resazurin ^25^. To basal medium trace elements and vitamins were added, for each trace element solution 1 mL L^-1^ and for the vitamin solution 0.2 mL L^-1^. The acid trace element solution consisted of (mM): H_3_Bo_4_, 1; CoCl_2_, 0.5; CuCl_2_, 0.1; FeCl_2_, 7.5; NiCl_2_, 0.1; MnCl_2_, 0.5; ZnCl_2_, 0.5 and HCl, 50. The alkaline trace element solution consisted of (mM): Na_2_MoO_4_, 0.1; Na_2_SeO_3_, 0.1; Na_2_WO_4_, 0.1 and NaOH, 50. The vitamin solution consisted of (g L^-1^): biotin, 0.02; cyanocobalamin, 0.1; niacin, 0.2; *p*-aminobenzoic acid, 0.1; pantothenic acid, 0.1; pyridoxine, 0.5; riboflavin, 0.1 and thiamine, 0.2. Medium was supplemented with 5 mM glycerol, 20 mM sodium sulfate, 0.1 g yeast extract and 1.5 mM of L-cysteine hydrochloride as reducing agent. All compounds were autoclaved except the vitamins, which were filter-sterilized through a 0.22 μM pore size polyethersulfone filter (Advanced Microdevices, Tepla, India). The reactors and 20 L medium tanks were sparged overnight and glycerol, vitamins and L-cysteine were added aseptically from sterile stock solutions.

### Reactor operation

Reactor cultivations were performed in 1.3 L DASGIP Parallel Bioreactors (Eppendorf, Hamburg, Germany) with a working volume of 0.75 L. Temperature was maintained at 30°C and an agitation rate of 100 rpm was ensured with overhead drives and two Rushton-type impellers at different heights. Reactors were also equipped with a redox probe (Hamilton, Reno, NV, USA). The pH was controlled with a pH probe and adjusted by automatic addition of 0.5 M or 1 M HCl. To maintain anoxic conditions and prevent sulfide accumulation the reactors were continuously sparged with N_2_/CO_2_ (80/20 v/v%) at a rate of 6 L h^-1^. Throughout the cultivations medium tanks were sparged with N_2_. A mid-exponential phase culture of *A. acetoxydans* was used to inoculate each reactor (inoculum 2% v/v). Growth was monitored off-line by optical density at 600 nm (OD_600_) using a Shimadzu UV-1800 spectrophotometer (Shimadzu, Kyoto, Japan). From the OD_600_ the cell dry weight (𝐶_𝐷w_) was calculated according to the ratio 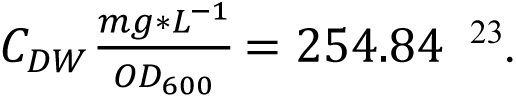 Reactor cultivations started as batch and in early exponential phase switched to continuous chemostat cultivation by turning on the feed and effluent pumps. Throughout the cultivations a dilution rate of 0.015 h^-1^ was used unless stated otherwise. Steady state was defined as less than 15% variation in OD_600_ and main extracellular metabolite concentrations (sulfate and glycerol) over three hydraulic retention times (HRT). Cultivations were terminated for technical reasons, not culture collapse.

### Metabolic activity assays

Metabolic activity assays were performed in 117 mL microcosm cultivation flasks with a 1.5 atm N_2_/CO_2_ (80:20, v/v) headspace, Flasks were sealed with butyl rubber stoppers (Rubber BV, Hilversum, Netherlands) and aluminum crimps. Duplicate microcosms were inoculated with 30 mL of culture from reactor R1 at the end of the cultivation. The glycerol concentration in R1 was increased to 2 mM and by automatic HCl addition the pH in the reactor, starting from pH 2.9, was lowered to 2.7, 2.5 and 2.3. At each pH two microcosms were supplemented with 30 mL culture and incubated statically at 30°C. Samples were taken at the tart off the incubation and after 1, 3 and 7 days for pH, OD_600_ and extracellular metabolite analysis.

### Analytical methods

Throughout the reactor runs samples were taken to monitor activity. For all measured compounds, the detection limit was approximately 100 µM. High performance liquid chromatography (HPLC) was used to measure glycerol and acetate on a Shimadzu LC2030c plus (Shimadzu, Kyoto, Japan) equipped with a Shodex SH1821 column (Shodex, Kyoto, Japan) and a differential refractive index detector Shimadzu RID-20A (Shimadzu, Kyoto, Japan). As eluent 5 mM sulfuric acid was used at 0.8 mL min^-1^ and the column was kept at 45°C. L-cysteine was measured after a derivatization pre-treatment with dabsyl chloride dissolved in acetonitrile. Supernatant of derivatized samples was measured on a Shimadzu LC2030c plus (Shimadzu, Kyoto, Japan) equipped with a Poroshell 120EC-C18 (Agilent, Santa Clara, CA, USA) and a differential refractive index detector Shimadzu RID-20A (Shimadzu, Kyoto, Japan). Sulfate was measured by anion exchange chromatography on a Dionex ICS-2100 (Dionex, Sunnyvale, CA, USA) equipped with a Dionex IonPac AS19 column (Dionex, Sunnyvale, CA, USA) operated at 30°C and a suppressed conductivity detector. As eluent KOH solution with a concentration gradient ranging from 10 to 40 mM was used at a flow rate of 0.4 mL min^−1^.

For elemental composition determination 10 mL samples were taken and acidified with HNO_3_ (1% v/v) and stored at 4°C until analysis. The metal elemental composition was determined in triplicate using an Avio 500 inductively coupled plasma-optical emission spectrometry (ICP-OES) (PerkinElmer, Waltham, MA, USA). Calibration standards were prepared from single elements and acidified with HNO_3_ (1% v/v), the calibration lines ranged from 20-100 µg L^-1^, 200-1000 µg L^-1^ and 2-10 mg L^-1^ depending on the measured elements. The results were automatically analyzed using Syngistix (PerkinElmer, Waltham, MA, USA).

### Transmission electron microscopy

Cell morphology and size was investigated by transmission electron microscopy (TEM). For the control samples, cells were harvested in mid-exponential phase at pH 5.0, end samples were taken 76 days into the cultivation with R1, R2 and R3 at pH 2.9 (approaching steady state), 2.9 (washing out) and pH 3.2 (approaching steady state). For each sample a 2 mL aliquot was centrifuged at 10000 x *g* for 5 minutes and the pellet resuspended and incubated for 1 hour at room temperature in fixative solution (2.5% glutaraldehyde, 2% paraformaldehyde and 0.1 M phosphate citrate buffer). The samples were prepared using gelatine and dehydrated with a graded series of ethanol as previously described ^23^. Dehydrated samples were embedded in Spurr epoxy resin ^26^ and cut into ultrathin sections with a Leica EM UC7 ultramicrotome (Leica, Wetzlar, Germany). The sections were poststained with uranyl-acetate and lead citrate and imaged using a JEOL JEM-1400 series 120 kV TEM (JEOL, Tokyo, Japan). From the TEM micrographs the cell sizes were obtained.

### Nucleic acid extraction

At steady state, 60 mL was sampled from the reactor and directly added to 100 mL of ice cold sterile reduced medium, followed by centrifugation at 10000 x *g* for 10 min at 4°C (ThermoFisher, Waltham, MA, USA). Obtained pellets were resuspended in 10 mL TE buffer and transferred to sterile 50 mL tubes (Greiner, Frickenhausen, Germany). A 2 mL aliquot was taken for DNA extraction and the remainder was centrifuged at 10000 x *g* for 10 min at 4°C. Supernatant was discarded and obtained pellets were stored at-80°C. RNA isolation was done as described previously ^27^. For DNA extraction, 2 mL of reactor sample was centrifuged at 10000 x *g* for 10 min at 4°C. The MasterPure Gram Positive DNA Purification kit (Epicentre, Madison, WI, USA) was used according to the manufacturer’s instructions. DNA samples were sent to Novogene (Novogene, Cambridge, UK) and sequenced using Illumina HiSeq (Illumina, San Diego, CA, USA) resulting in paired-end reads of 150 bp.

### Transcriptome and genome analysis

Raw sequence data was trimmed for Illumina adapters and paired using Trimmomatic 0.38 ^28^ with the following settings: SLIDINGWINDOW: 4 bases, quality ≥20, average read score >5, MINLEN: 100. Quality trimmed paired-end reads were checked using FastQC 0.11.9 and MultiQC 1.11 ^29,30^. Reads were mapped with BWA-MEM 0.7.17 ^31^ against the reference genome of *A. acetoxydans* (Genbank accession number: LR746496). Before mapping 66 structural RNA (rRNA, sRNA and tRNA) genes were removed from the dataset. Count matrices with mapped reads were obtained using featureCounts 2.0.1 with paired-end reads enabled ^32^ and mapping quality was assessed with SAMtools (flagstat) 2.0.4 ^33^. To test for significant statistical differences in gene expression profiles, read counts were analyzed with the R package DESeq2 1.30.1 ^34^. Differentially expressed genes were identified as genes with an expression change of at least twofold and an adjusted p-value ≤0.05. For genome analysis the quality of raw sequence data and absence of Illumina sequencing adapters was checked using FastQC 0.11.9 and MultiQC 1.11 ^29,30^. Obtained raw reads were compared to the closed genome of the parental genome of *A. acetoxydans* DSM 29876^T^ (Genbank accession number: LR746496) using breseq 0.35.5 with default settings ^35^. Functional categories were used for general analysis and were defined by assigning *A. acetoxydans* genes to clusters of orthologous groups (COGs) using eggNOG 5.0 ^36,37^.

### Lipid analysis

At the same time as the transcriptome samples, 20 mL sample of each reactor was taken and centrifuged at 10000 x *g* for 10 min at 4°C. Obtained pellets were resuspended in phosphate buffered saline solution (0.01 M phosphate, 0.27 mM KCl, 0.137 M NaCl pH 7.4) and again centrifuged at 10000 x *g* for 10 min at 4°C, the pellets were stored overnight at-20°C. Afterwards samples were lyophilized and stored in the dark at 4°C until further analysis. Freeze-dried biomass was subjected to acid hydrolysis followed by derivatization and detection by gas chromatography-mass spectrometry to determine the composition of core lipids ^38^.

## Results

### *Acididesulfobacillus acetoxydans* is metabolically active at pH values representing AMD acidity

To determine the physiological limits of *A. acetoxydans* under proton stress, continuous anaerobic cultivations were performed across a controlled pH gradient ranging from its optimum (pH 5.0) to values characteristic of acid mine drainage (pH 2.9). *A. acetoxydans* was grown anaerobically in three independent chemostat cultures (R1-R3). At pH 5.0, steady states reached an OD_600_ of 0.55 ± 0.03 (average ± standard deviation) after 3 hydraulic retention times (HRT, ∼66 hours each). Successive pH decreases to 4.4, 3.5 and 3.2 yielded steady states with OD_600_ values of 0.45 ± 0.02, 0.46 ± 0.02 and 0.47 ± 0.03, respectively (R1-R2, Figure 1). A shift from pH 3.5 to 2.9 resulted in biomass washout, but recovery occurred after readjustment to pH 3.2, indicating reversible inhibition rather than loss of viability. After steady state at pH 3.2, re-lowering the pH to 2.9 with a reduced dilution rate (0.010 h^-1^) resulted in measurable growth in both R1 and R2 (Table S2), with growth rates of 0.0095 h^-1^ in R1 and 0.0081 h^-1^ in R2 before partial washout. Lower pH values might have been attainable, but this was not tested as the reactors were terminated at pH 2.9. When growing at pH 2.9, representing a 126-fold increase in proton concentration compared to its optimum, *A. acetoxydans* maintained continuous planktonic growth without biofilm formation or the development of protective microniches (*e.g.*, granulation). Routine light microscopy performed during steady state conditions revealed no morphological changes in *A. acetoxydans*. To verify this and potentially visualize changes in cell size or membrane thickness, transmission electron microscopy (TEM) samples were collected from each reactor at its final pH (day 76) and from a control flask culture at pH 5.0 (mid-exponential phase, also used as reactor inoculum). TEM micrographs confirmed uniform morphology and no change in cell size or aggregation across pH conditions (Table S3; images available at Figshare DOI:10.6084/m9.figshare.30407224). Follow-up microcosm activity assays from R1 derived biomass at pH 2.9 demonstrated metabolic activity at pH 2.9, 2.7 and pH 2.5 (Figure S1), whereas no activity occurred at pH 2.3. This expands the known physiological range for this species beyond the previously reported threshold of pH 3.8 and aligns with the typical acidity of AMD (pH 2.2-2.8) ^23^.

**Figure 1.**
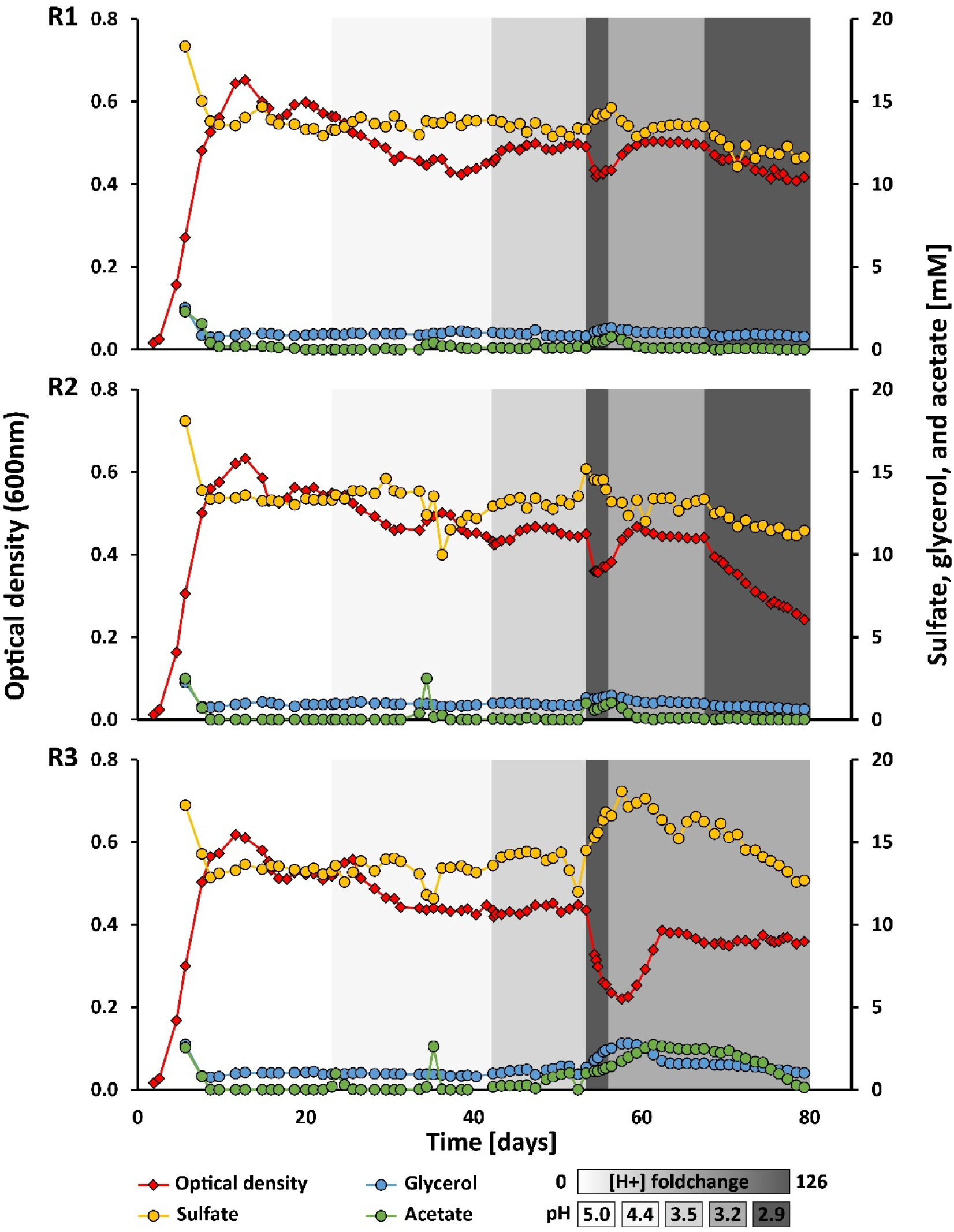
Optical density (OD_600_) and glycerol, acetate and sulfate concentrations of *A. acetoxydans* grown in triplicate pH-controlled bioreactors (R1-R3). The initial metabolite measurement marks the switch from batch to continuous cultivations with a dilution rate of 0.015 h^-1^. Grayscale intensity represents the pH and respective change in proton concentration compared to pH 5.0. The dilution rate in R1 and R2 at pH 2.9 was lowered from 0.015 to 0.010 h^-1^.

### Robust sulfate reduction rates at decreasing pH and increased impact of metal toxicity

To evaluate how acid stress affected energy metabolism and reactor performance, sulfate reduction and carbon oxidation were quantified across decreasing pH values. At each tested pH *A. acetoxydans* coupled growth to sulfate reduction and complete glycerol oxidation. The biomass specific sulfate reduction rate (qS, µmol mg CDW^-1^ day^-1^) of R1-R3 was 22.8 ± 1.4 at pH 5.0 and remained stable at decreasing pH until pH 3.2 and even pH 2.9 (Table S4). Although sulfate was in excess, approximately 1 mM of glycerol persisted in the reactors. Throughout the cultivations, glycerol was completely oxidized to carbon dioxide under steady state conditions. Temporary acetic acid accumulation occurred only during operational perturbations in the system (*e.g.* slipping of pumps), or the decrease to pH 2.9, after readjustment to pH 3.2 activity the full oxidation of glycerol resumed.

Reactor performance differences at low pH correlated with the metal leakage from affected stainless-steel reactor components (Table S5). Metal and major element concentrations were measured in samples collected on days 73, 75 and 80. A metal leakage ratio of 1:2:3 was observed for R1, R2 and R3, respectively. At day 80 of the cultivations, these elevated concentrations of stainless-steel components were iron: 16, 38, 73 mg L^-1^; chromium: 4, 8, 17 mg L^-1^; nickel: 2, 5, 11 mg L^-1^; and manganese: 293, 649, 976 µg L^-1^ for R1, R2 and R3, respectively. These rising concentrations coincided with a decline in biomass but stable volumetric sulfate reduction rates, particularly in R1 and R2 at pH 2.9, implying increased cell-specific activity to handle metal and proton stress, most likely due to increased maintenance energy expenditure due to proton leakage and potential metal toxicity.

### Genome analysis revealed no evidence of laboratory evolution

To determine whether the observed pH tolerance resulted from physiological adaptation or genetic change, whole-genome sequencing was performed. Using the published (closed) genome of *A. acetoxydans* (NCBI accession number: SAMEA6497484) as reference, sequenced DNA reads from R1 at pH 5.0 were used to generate a single nucleotide polymorphism (SNP) fingerprint and determine mutations. The fingerprint was compared to the reads obtained from R1-R2 (pH 2.9) and R3 (pH 3.2) at day 79. This ensured that only gained mutations during the experiment were identified, including gene gain or gene loss events. Relative to the reference, no gene losses or acquisitions were detected, and each condition contained the same 19 SNPs (Table S6). A single additional SNP was observed only in R1 at pH 5.0, consisting of an extra cytosine insertion in gene DEACI_0987, which encodes a hypothetical protein with no predicted physiological function. These results confirm that the observed pH tolerance reflects physiological adaptation rather than laboratory evolution.

### Transcriptome response to low pH revealed globally reduced gene expression

To identify the molecular basis of the observed stability under proton stress, transcriptomic profiles were compared for reactors operated at pH 5.0, 3.5, 3.2 (R1–R2) and 2.9 (R1–R2). The largest transcriptomic shift occurred between pH 5.0 and the lower pH conditions, whereas profiles for pH 3.5, 3.2 and 2.9 were nearly identical (Table 1). Principal component analysis confirmed that variance was primarily driven by separation of pH 5.0 from the acidic conditions and that biological replicates from different reactors clustered together (Figure S2). Mapped reads were converted to transcripts per million (TPM); expression levels of genes with <10 TPM were considered negligible, 10-50 TPM low, 50-300 TPM medium and >300 TPM high. The total number of more low, medium or highly expressed genes (≥10 TPM) decreased with decreasing pH (Table S7), indicating that *A. acetoxydans* expressed a smaller fraction of its genome under more acidic conditions. Differentially expressed genes (DEGs), classified by clusters of orthologous groups, were most abundant in cell-wall and membrane biogenesis (COG M), amino-acid metabolism (E) and inorganic-ion transport (P), interestingly cell-motility genes (N) were consistently downregulated at decreasing pH (Table S8).

**Table 1.**
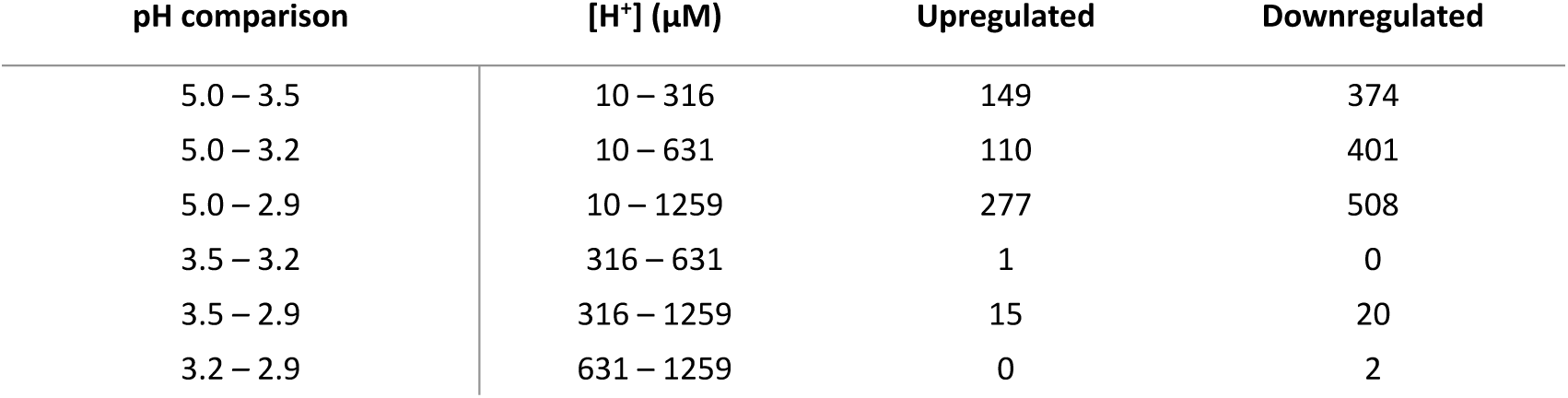
Number of differentially expressed genes (DEGs) between pH conditions including absolute proton concentrations [H^+^], *A. acetoxydans* has 4104 protein-encoding genes. The gene-expression profiles at pH 3.2 and 2.9 are derived from reactors R1 and R2.

| pH comparison | $[H^+]$ ($\mu M$) | Upregulated | Downregulated |
| --- | --- | --- | --- |
| 5.0 – 3.5 | 10 – 316 | 149 | 374 |
| 5.0 – 3.2 | 10 – 631 | 110 | 401 |
| 5.0 – 2.9 | 10 – 1259 | 277 | 508 |
| 3.5 – 3.2 | 316 – 631 | 1 | 0 |
| 3.5 – 2.9 | 316 – 1259 | 15 | 20 |
| 3.2 – 2.9 | 631 – 1259 | 0 | 2 |

### Central metabolism remained stable, while expression of motility and sporulation related genes decreased at low pH

Consistent with the physiological data, no major changes were observed in central energy or carbon metabolism, confirming metabolic stability across the different pH values. In the central carbon pathway, glyceraldehyde-3-phosphate dehydrogenase (DEACI_2591) and pyruvate kinase (DEACI_2592) were upregulated at lower pH compared to pH 5.0, supporting the increased relative importance of preventing acetic acid accumulation at lower pH. In contrast, genes linked to energy demanding processes such as flagellar biosynthesis and sporulation were consistently downregulated at lower pH (Tables S9-S10).

### Potassium transport encoding genes are upregulated at low pH

Genes encoding for proteins involved in proton and ion homeostasis were expressed under all tested conditions (Table S11). Technically, the F_O_F_1_-ATPase (*atpA–I*) and the flagellum-specific ATPase (*fliL*) could export protons at the cost at ATP; their genes were expressed at medium to high levels in all conditions, indicating constitutive expression throughout the pH range. No DEGs were detected for active proton export or proton antiporters. The high-affinity K^+^-transporting ATPase (*kdpABC*, DEACI_3037-3039) showed consistent upregulation at low pH, suggesting that maintenance of intracellular K⁺ pool has increased relative importance in countering proton influx at lower pH. The associated regulatory histidine kinase pair (*kdpDE*, DEACI_1068–1069) did not change in expression. Other cation transporters remained transcriptionally constant. Combined this indicates an emphasis on maintaining a chemiosmotic gradient over increased active proton export at lower pH.

### Branched chain amino acid biosynthesis pathways are upregulated at low pH

Several genes related to amino acid metabolism and transport changed in expression under acidic conditions (Table S12). The branched chain amino acid (BCAA) transporter *livFGHMJ* (DEACI_0431–0435) was downregulated, whereas multiple BCAA biosynthetic genes (*ilv-leu* cluster, DEACI_2327–2336) were upregulated, suggesting a shift in amino acid synthesis. Similarly, proline synthase (*proABC*, DEACI_2337-2339) and most genes in the tryptophan biosynthesis pathway (*trpABDGE*, DEACI_2714-2718) were upregulated at low pH although the genes encoding the final tryptophan biosynthesis proteins *trpCF* (DEACI_2743-2744) were expressed at medium levels in all conditions. *A. acetoxydans* encodes two glutamine synthetases (*glnA1-2*, DEACI_1897, 1898), both were expressed in all conditions. Their transcription is regulated by the nitrogen regulatory protein P-II (*glnB*), for which two of several gene copies (DEACI_1382, 2355) were expressed. At low pH, both *glnA2* and *glnB1* were upregulated, suggesting nitrogen assimilation towards amino acid synthesis for stress adaptation and possible cytoplasmic buffering through ammonium incorporation.

### DNA repair and metal homeostasis genes showed limited transcriptional change

Genes involved in DNA repair, protein quality control, and oxidative stress response were constitutively expressed across all pH conditions (Table S13). The chaperonin encoding genes *groES* (DEACI_1389, 2469) and *groEL* (DEACI_1390) were among the most highly expressed genes throughout the experiment. Despite decreasing pH, only minor transcriptional adjustments occurred: at pH 2.9, the UvrABC excision-repair genes (*uvrAB*, DEACI_0322– 0323) were upregulated, whereas *uvrC* remained constant. Among peroxiredoxins, DEACI_0888–0889 were downregulated and DEACI_2586 upregulated, reflecting a limited oxidative stress response. As Fe, Cr, Ni, and Mn accumulated in reactor effluents, the metal-uptake permease (*zupT*, DEACI_0143) and ferritin (DEACI_0071) were downregulated relative to pH 5.0, while the nickel transporter (*hoxN*, DEACI_1037) and cobalt/nickel transporter operon (*cbiQMO*, DEACI_2205, DEACI_2207-2208) were upregulated (Table S14). Other metal-resistance genes remained transcriptionally stable. Overall, the transcriptional response of *A. acetoxydans* to increasing proton concentrations was modest, characterized by reduced expression profiles, minimizing energetically demanding processes such as motility and amino acid transport with selective upregulation of genes involved in ion transport and repair mechanisms. This suggests that adaptation to low pH primarily involves structural rather than transcriptional adjustments, such as membrane remodeling

### Membrane lipid composition changes progressively with decreasing pH

Analysis of the hydrolysis-derived lipids (core lipids) of *A. acetoxydans* revealed clear compositional remodeling with decreasing pH. Detected lipids include fatty acids, hydroxy fatty acids, dimethylacetals (DMAs; formed upon acid hydrolysis of plasmalogens, or also known as alkenyl ethers) and mono alkyl glycerol ethers (MGEs; derived from non-plasmalogen ether lipids, also known as alkyl ethers). While the overall lipid classes were consistent there was a slight increase in the already high percentage of saturated lipids at decreasing pH and the proportion with methyl moieties (9-and 10-methyl) increased substantially (Table 2). At the same time, the relative abundance of *iso*-and *anteiso*-fatty acids declined from 64% to 45%, whereas 9-and 10-methyl-substituted *iso*-an *anteiso*-fatty acids increased from 23% to 38% toward pH 2.9. This trend, together with the high proportion of saturated lipids, indicates significant membrane remodeling at low pH.

**Table 2.** Simplified overview of the relative abundance (% of total, average ± standard deviation) of hydrolysis-derived lipids of *A. acetoxydans* at each pH, the complete table can be found in Table S15. The detected core lipids included fatty acids (FAs), β-OH FAs, dimethylacetals (DMAs; formed upon acid hydrolysis of plasmalogens) and mono alkyl glycerol ethers (MGEs). For each of the categories the sum of ethers, straight chain fatty acids, branched fatty acids (*iso*, *i.* and *anteiso*, *a.i.*) and unsaturated fatty acids is depicted. Here fatty acids are named as Cx:y with X being the number of carbons and y the number of double bonds, *i: iso* and Me: methyl group. The asterisk (*) depicts unknown methyl location for the dialkyl glycerol ether (DGE) and therefore has unknown priming precursor. The priming precursors consist of acetate (acetyl-CoA), leucine (3-methylbutyryl-CoA), isoleucine (2-methylbutyryl-CoA) and valine (2-methylpropanyl-CoA).

| Compound |  | precursor | pH 5.0<br>(R1-R3) | pH 4.4<br>(R1-R3) | pH 3.5<br>(R1-R3) | pH 3.2<br>(R1-R3) | pH 2.9<br>(R1-R2) |
| --- | --- | --- | --- | --- | --- | --- | --- |
| Fatty acids | *i.* C15:0 | leucine | 29.4 ± 12.4 | 26.0 ± 7.3 | 28.0 ± 4.0 | 14.9 ± 5.9 | 13.4 ± 14.8 |
|  | *i.* C16:0 | valine | 1.1 ± 0.5 | 5.8 ± 2.4 | 6.6 ± 1.4 | 10.7 ± 1.6 | 10.1 ± 3.3 |
|  | C16:0 | acetate | 2.8 ± 0.9 | 1.8 ± 2.3 | 2.2 ± 1.6 | 1.8 ± 0.9 | 2.0 ± 0.8 |
|  | 10-Me *i.* C16:0 | isoleucine | 0.4 ± 0.7 | 5.1 ± 2.5 | 6.2 ± 0.5 | 8.8 ± 1.0 | 10.8 ± 0.9 |
|  | *i.* C17:0 | leucine | 2.2 ± 0.3 | 1.6 ± 0.8 | 1.6 ± 0.2 | 1.3 ± 0.3 | 1.2 ± 0.0 |
|  | 10-Me *i.* C17:0 | leucine | 4.6 ± 1.5 | 4.8 ± 2.6 | 3.9 ± 0.3 | 3.0 ± 0.2 | 3.4 ± 0.7 |
|  | Sum |  | **40.5 ± 11.2** | **45.1 ± 5.8** | **48.5 ± 2.7** | **40.5 ± 6.5** | **40.9 ± 15.7** |
| DMAs | *i.* C15:0 | leucine | 4.9 ± 0.5 | 2.1 ± 1.0 | 1.3 ± 0.8 | 0.3 ± 0.0 | - ± - |
|  | *i.* C16:0 | valine | 2.6 ± 0.2 | 7.3 ± 3.3 | 6.6 ± 2.7 | 6.8 ± 1.8 | 4.7 ± 3.3 |
|  | *i.* C17:0 | leucine | 5.2 ± 1.9 | 2.6 ± 0.6 | 2.5 ± 1.1 | 1.4 ± 0.4 | 0.8 ± 0.4 |
|  | 10-Me *i.* C17:0 | leucine | 3.3 ± 1.0 | 1.7 ± 0.5 | 1.0 ± 0.5 | 0.6 ± 0.3 | 0.6 ± 0.2 |
|  | Sum |  | **15.9 ± 2.5** | **13.7 ± 3.4** | **11.4 ± 4.9** | **9.2 ± 2.5** | **6.1 ± 3.9** |
| β-OH<br>FAs | Sum |  | **2.5 ± 0.1** | **2.1 ± 0.9** | **2.1 ± 0.4** | **3.8 ± 2.7** | **3.7 ± 1.8** |
| MGEs | *i.* C15:0 | leucine | 13.4 ± 3.4 | 5.4 ± 2.6 | 2.8 ± 0.5 | 1.6 ± 0.1 | 1.5 ± 0.5 |
|  | *i.* C16:0 | valine | 1.9 ± 0.5 | 5.5 ± 0.2 | 6.8 ± 0.9 | 11.6 ± 0.8 | 12.1 ± 4.5 |
|  | C16:0 | acetate | 0.2 ± 0.3 | 0.9 ± 1.6 | 2.7 ± 1.7 | 3.4 ± 1.8 | 1.2 ± 1.7 |
|  | 10-Me *i.* C16:0 | valine | 3.1 ± 1.7 | 6.5 ± 2.6 | 6.4 ± 2.6 | 9.6 ± 4.1 | 16.8 ± 6.5 |
|  | 10-Me C16:0 | acetate | 0.3 ± 0.2 | 1.6 ± 1.2 | 1.9 ± 2.2 | 2.4 ± 2.0 | 0.9 ± 0.5 |
|  | *i.* C17:0 | leucine | 2.8 ± 0.3 | 2.4 ± 1.6 | 2.2 ± 0.2 | 1.9 ± 0.3 | 1.5 ± 0.4 |
|  | 10-Me *i.* C17:0 | leucine | 11.4 ± 3.5 | 7.7 ± 4.2 | 6.6 ± 0.9 | 5.2 ± 1.4 | 6.4 ± 2.4 |
|  | Sum |  | **33.0 ± 8.2** | **30.0 ± 7.4** | **29.3 ± 2.1** | **35.6 ± 2.0** | **40.4 ± 16.5** |
| DGEs | C30:0* |  | 0.6 ± 0.2 |  |  |  |  |
| Sum hydrolysis-derived lipids | Ether |  | 49.6 ± 9.3 | 43.6 ± 5.1 | 40.7 ± 3.0 | 44.8 ± 4.5 | 46.5 ± 12.6 |
|  | Unsaturated |  | 15.9 ± 2.5 | 13.7 ± 3.4 | 11.4 ± 4.9 | 9.2 ± 2.5 | 6.1 ± 3.9 |
|  | Straight chain |  | 2.9 ± 1.1 | 2.7 ± 3.8 | 4.9 ± 3.3 | 5.2 ± 2.7 | 3.2 ± 2.5 |
|  | *i.* & *a.i.* |  | 64.2 ± 9.2 | 58.7 ± 7.8 | 58.3 ± 7.2 | 50.5 ± 6.7 | 45.3 ± 16.4 |
|  | 10-Me straight chain |  | 0.3 ± 0.2 | 1.6 ± 1.2 | 1.9 ± 2.2 | 2.4 ± 2.0 | 0.9 ± 0.5 |
|  | 9- + 10-Me *i.* & *a.i.* |  | 22.7 ± 6.0 | 25.7 ± 4.3 | 24.1 ± 3.5 | 27.2 ± 6.6 | 38.0 ± 10.3 |

The biosynthesis of core lipids is initiated by specific priming precursors: acetate (acetyl-CoA), leucine (3-methylbutyryl-CoA), isoleucine (2-methylbutyryl-CoA), and valine (2-methylpropanyl-CoA). These priming precursors determine whether the resulting lipid is straight-chain, *iso*-, or *anteiso*-branched. Analysis of the precursor distribution showed a major shift across the tested pH range: valine increased from 10.2% to 49.4%, leucine decreased from 82.3% to 30.8%, and isoleucine increased from 0.8% to 11.2% between pH 5.0 and 2.9 (Table S15). Despite this pronounced remodeling, genes directly involved in fatty acid synthesis remained largely unchanged except for *ilvE*, which encodes the branched-chain amino-acid transaminase responsible for converting leucine (Table S16). This coordinated remodeling of core lipids, shift in priming precursor and upregulation of BCAA metabolism indicates a major structural and physiological response to increasing proton stress. The shift from leucine (C_5_) to valine (C_4_) primers shortens the average chain length and increases branching density, reinforcing membrane tightness under acid stress and by increasing mid-chain methyl branching, the organism effectively tunes rigidity and fluidity to withstand AMD-like acidity.

## Discussion

Continuous cultivations demonstrated that the moderate acidophile *A. acetoxydans* can grow planktonically and remain metabolically active down to pH 2.9. Subsequent microcosm cultivations demonstrated metabolic activity at pH 2.5. This extends the known lower limit for acidophilic sulfate-reducing bacteria (aSRB) and closes the gap between laboratory isolates as the same pH (2.2-2.8 range) is typical of many AMD environments, including ones with reported biosulfidogenesis ^5,6,19,39–41^. The previously reported growth limit of pH 3.8 coincides with the pH of the upper sediment layer of Rio Tinto (Huelva, Spain), a well-studied AMD environment, whereas the identified limit here matches, or even exceeds, the pH observed throughout both sediment and water column ^42^. For a long time the accepted view was the SRB prefer a circumneutral environment, with observations of SRB at low pH values being due to microniches ^43,44^. These results demonstrate that *A. acetoxydans* can maintain both growth and sulfate reduction under conditions previously thought incompatible with SRB activity and provide the first controlled physiological evidence of planktonic aSRB activity at pH values characteristic of AMD waters.

Triplicate chemostats yielded stable steady states across the tested pH range. The drop in optical density after switching from pH 3.5 to pH 2.9 demonstrated that gradual pH adaptation is required for continuous cultivation of aSRB at increasingly acidic conditions. Under excessive stress, aSRB may sporulate, hampering cultivation. Sulfate reduction rates were stable across the different pH values; when expressing as volumetric sulfide production rates (VSPR) they remained between 105-63 mg L^-1^ day^-1^ (Table S4). These values are comparable to those reported for mixed or acclimated sulfidogenic consortia operating at similar pH values (Table S17). Direct comparisons to other systems should be made cautiously, as most reference reactors operated with high biomass sludge or granular matrices that provide protective microniches. In contrast, the continuous stirred tank reactors used in this study contained planktonic cells and were optimized for physiological rather than process performance. At pH 2.9, sulfate reduction was maintained even when growth rates fell below the imposed dilution rate. Metal release from stainless-steel components (Fe, Cr, Ni, and Mn) likely contributed to additional energy burden through metal detoxification and imposed stress. Despite this, VSPR remained relatively stable, underscoring the robustness of aSRB for biosulfidogenesis at low pH.

The persistent glycerol surplus of approximately 1 mM across all reactor cultivations indicates that the system was not energy-substrate limited. It is unclear why this residual glycerol remains, as it cannot be attributed to a shortage in vitamins or trace elements. All glycerol including the additional glycerol supplied up to 2 mM, was completely oxidized in subsequent microcosm incubations. Acetic acid (p𝐾_𝑎_ = 4.76) becomes increasingly toxic to microorganisms at low pH as the proportion of its undissociated form rises under acidic conditions. This neutral form can freely diffuse across the cell membrane, acidify cytoplasm and disrupt anion homeostasis ^24,45^. The residual glycerol at low pH likely reflects a shift in energy allocation toward complete oxidation preventing acetic acid toxicity, which only accumulated during temporary reactor perturbations and increased maintenance demand, rather than incomplete substrate utilization. Supporting this interpretation, there was no change in cell size which is often a characteristic of nutrient limitation ^46^.

No genomic changes were detected between conditions, excluding laboratory evolution as the cause of the observed tolerance. Transcriptome profiles showed only modest changes, main observations were decreased gene expression profiles and downregulation of genes related to motility at lower pH values, which is a known phenomenon for both nutrient limitation and acid stress ^46,47^. Acidophiles have several mechanisms to cope with low pH, which can be categorized into mechanisms impeding the protons from penetrating the cell and mechanisms that export or consume protons that enter the cell ^17,21,48^. In *A. acetoxydans*, no significant upregulation of proton exporting ATPases or proton consuming decarboxylation pathways was observed ^49^. The lack of enhanced expression of proton extrusion and consumption mechanisms indicate that pH homeostasis at lower pH mainly revolved around proton impeding mechanisms. To counter the large pH gradient (ΔpH) across the membrane, acidophiles generate a reversed membrane potential (Δψ, inside the cell positive) by the influx of cations creating a chemiosmotic gradient to repel proton entry ^17,50^. In particular the importance of potassium has been observed in a variety of acidophiles ^51,52^. The upregulation of genes encoding the K^+^-transporting ATPase KdpABC, suggests an increased importance at lower pH values and potentially confers resistance to higher metal concentrations ^53^. Multiple potassium transporting systems exist such as the Kdp, Kup, Kch and Trk systems ^21,48^. Interestingly, of the characterized aSRB (Table S1), only *Desulfosporosinus metallidurans* encodes TrkAH instead of KdpABC. In addition to KdpABC, *Desulfosporosinus acididurans* also encoded Kup. An interesting comparison would be on the evolution of these potassium systems and other acid resistance genes in aSRB as was done for the *Acidohalobacter* genus ^21^. No upregulation of genes encoding other cation transporters was observed suggesting that K⁺ import is the dominant mechanism to maintain a reversed membrane potential.

Genes involved in DNA, RNA and protein were expressed across all pH conditions, indicating a continuous investment rather than induced stress response. The chaperonin genes *groES* and *groEL* were among the most highly expressed under all conditions, suggesting the importance in refolding and stabilizing proteins exposed to proton and metal stress ^54^. Only limited changes in gene expression occurred at pH 2.9: the *uvrAB* components of the UvrABC nucleotide-excision-repair system were upregulated, whereas *uvrC* remained constant. This system repairs acid-induced DNA lesions, with *uvrA* mutants showing increased pH sensitivity ^55,56^. Moreover, *uvrA* is often upregulated when microorganisms are exposed to either acid or organic acid stress ^57,58^ with overexpression of *uvrA* enhancing acetic acid tolerance ^59^. The constitutive expression of these systems across tested pH suggests that *A. acetoxydans* continuously maintains repair and detoxification systems, which is likely essential yet does not differ noticeably at lower pH.

Branched-chain amino-acid (BCAA) synthesis, the *ilv-leu* gene cluster, was upregulated at low pH, indicating enhanced capacity for valine, leucine, and isoleucine biosynthesis. Although transcriptomics is unable to determine metabolic flux, core lipid priming precursor data revealed a pronounced shift from leucine toward valine and isoleucine derived priming precursors, suggesting altered metabolic routing toward shorter branched-chain precursors under acid stress. This synthesis shift could be an energy optimization mechanism as valine (C_4_) synthesis requires approximately 15% less energy than leucine (C_5_) synthesis as well as one less carbon ^46^. Both proline and tryptophan biosynthetic pathways were also upregulated. Intracellular proline accumulation has linked as acid stress response in the extreme acidophile *At. caldus* ^60^. Potentially proline acts as compatible solute to compensate for the increased influx of potassium, to maintain cytoplasmic solvent properties ^61^. Tryptophan synthesis is notable considering it is the second most expensive amino acid to synthesize ^46^. Further metabolomic analysis of intracellular amino acid pools and targeted supplementation or substrate selection at low pH could both clarify their specific contribution and serve as substrates to enhance aSRB in bioremediation.

Extreme acidophiles must maintain membranes with extremely low proton permeability ^17,62,63^. In *A. acetoxydans*, decreasing pH induced a shift toward more saturated lipids, which was also observed for the extreme acidophiles *Acidithiobacillus ferrooxidans* and *At. caldus* ^64,65^. This shift was observed previously in batch reactor cultivations in *A. acetoxydans* under acetic acid stress and comparing pH 3.9 to pH 5.0, indicating the importance of cultivation technique on membrane composition ^23,24^. The progressive increase in saturation at low pH values, indicates an increased rigidity at low pH to impede proton influx. Generally, *anteiso*-FAs promote a more fluid membrane structure than *iso*-FAs, as the methyl branch is further from the end of the fatty acid, which is beneficial at low pH ^66,67^. By extension, mid-branching may promote membrane fluidity to an even higher degree than *iso* or *anteiso* branching hence the 9-and 10-methyl mid-chain branching may enhance the fluidity of the lipid as has been suggested from molecular dynamics simulations ^68^. Such structural remodeling likely decreases energy costs related to membrane synthesis and ultimately in proton transport and repair, providing a physiological advantage for aSRB in AMD-like acidity.

This study addressed two key knowledge gaps: the lower pH tolerance of aSRB and the mechanisms underlying their activity under proton stress. The integrated physiological, transcriptomic, and lipidomic data show that aSRB tolerate extreme acidity through coordinated metabolic and structural adjustments rather than inducible stress responses. Together, these results demonstrate that *A. acetoxydans* retains metabolic activity through structural adaptation, integrating metabolic rerouting and lipid remodeling to withstand AMD-like acidity. Likely previous pH limits largely reflected cultivation constraints given the complexity of working with sporulating microorganisms. The tolerance range of *A. acetoxydans* and likely many other aSRB overlaps with the acidity of AMD waters and sediments, providing a physiological basis to interpret field observations of sulfate reduction in such environments. In engineered systems, this could reduce or eliminate the need for chemical neutralization and enable direct bioremediation of acidic mine effluents, underscoring the potential of aSRB for metal removal and selective recovery under extreme conditions. Future research should explore the interplay of proton and metal stress to guide the design of scalable AMD bioremediation technologies.

## Data Availability Statement

The datasets for this study have been deposited at the European Nucleotide Archive: https://www.ebi.ac.uk/ena. Raw DNA and RNA sequencing reads can be found under accession number PRJEB71105 and PRJEB71113, respectively. High-resolution transmission electron micrographs can be found on Figshare: 10.6084/m9.figshare.30407224.

## Acknowledgements

This work was financed by the Soehngen Institute for Anaerobic Microbiology Gravitation Program (SIAM 024.002.002), a Gravitation grant of the Netherlands Ministry of Education, Culture and Science. We like to thank Pieter Gremmen for his help on the elemental composition analysis, Anchelique Metz for core lipid workup and analysis, and Diana Sahonero Canavesi for the lipid discussions at the onset of this study.

## Supplementary Material

**Figure S1.** Overview pH profiles microcosm incubations and metabolite concentrations for pH 2.5.

**Figure S2.** Principal component analysis of obtained gene expression profiles.

**Table S1.** Chronological order of characterized acidophilic and acidotolerant SRB.

**Table S2.** Calculated growth rate of R1 and R2 after switching to pH 2.9.

**Table S3.** Overview of cell sizes obtained from selected TEM micrographs of *A. acetoxydans* at final pH of each chemostat compared to optimum pH.

**Table S4.** Calculated sulfate reduction rates for reactors at different obtained pH.

**Table S5.** Elemental composition analysis of both medium and influent of the different bioreactors.

**Table S6.** Overview of obtained single nucleotide polymorphism (SNP) fingerprint of *A. acetoxydans*.

**Table S7.** Overview of locus tags of *A. acetoxydans* and corresponding expression levels (transcripts per million) and DESeq2 statistics for the different conditions.

**Table S8.** Change in differentially expressed genes for each cluster of orthologous groups per pairwise pH comparison.

**Table S9-S14.** Annotation and expression of genes related to motility (**Table S9**), sporulation (**Table S10**), pH homeostasis (**Table S11**), amino acid metabolism and transport (**Table S12**), repair mechanisms (**Table S13**) and metal resistance (**Table S14**).

**Table S15.** Overview of the relative abundance (% of total) of fatty acids and ether lipids and corresponding priming precursors at different pH.

**Table S16.** Annotation and expression of genes related to lipid synthesis.

**Table S17.** Overview of high-rate sulfate-reducing sulfidogenic bioreactors operated under acidic conditions.

