## Supplementary material for "Proton stress adaptation in acidophilic sulfate-reducing bacteria: insights from *Acididesulfobacillus acetoxydans* for acid mine drainage bioremediation": Figure S1

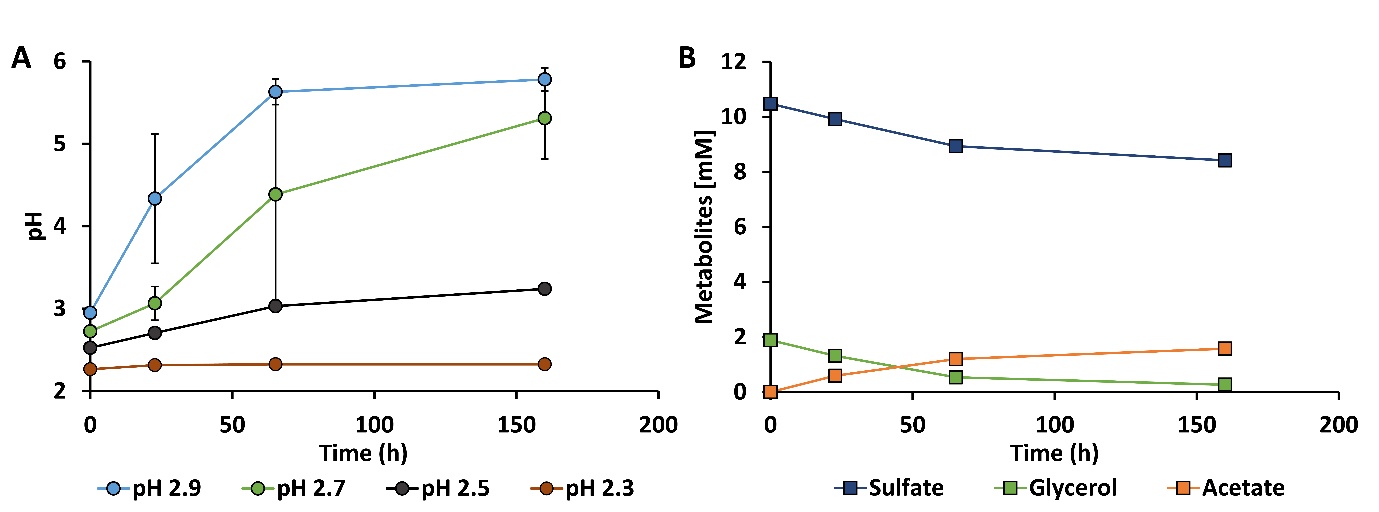
**Figure S1. (A)** pH profiles of (duplicate) microcosm incubations inoculated from reactor R1 at initial pH 2.9 supplemented with 2 mM of glycerol. (**B)** Concentration changes of sulfate, glycerol and acetate during incubations at pH 2.5, showing incomplete glycerol oxidation to acetate and confirming metabolic activity of *A. acetoxydans* at pH 2.5. Error bars indicate standard deviations.
