## Supplementary material for "Proton stress adaptation in acidophilic sulfate-reducing bacteria: insights from *Acididesulfobacillus acetoxydans* for acid mine drainage bioremediation": Figure S2

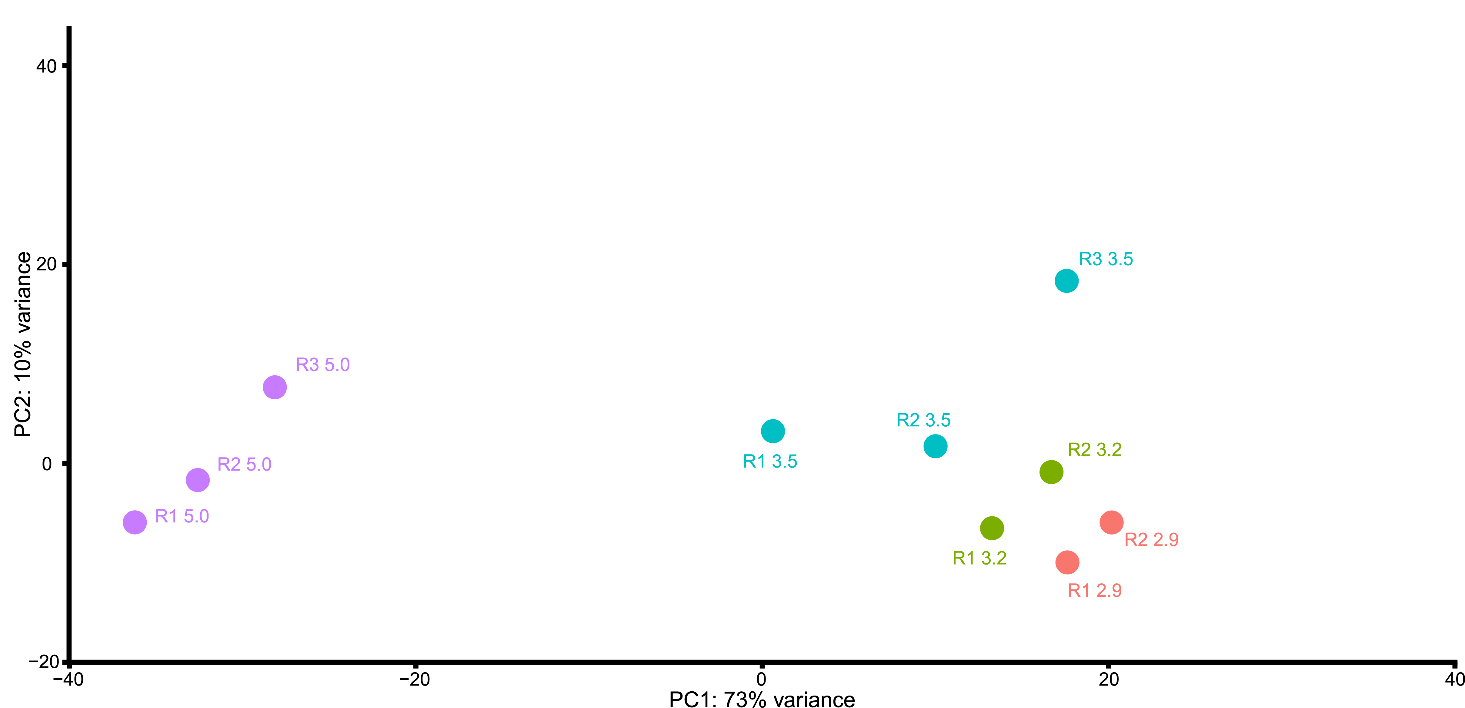
**Figure S2.** Principal component analysis of obtained gene expression profiles. R1, R2 and R3 depict the three bioreactors at different pH.
