## Supplementary material for "Proton stress adaptation in acidophilic sulfate-reducing bacteria: insights from *Acididesulfobacillus acetoxydans* for acid mine drainage bioremediation": Table S1

**Table S1.** Chronological order of characterized acidophilic and acidotolerant SRB, with their temperature and pH ranges including optimum, complete or incomplete organic acid oxidation and their origin. **Desulfothermobacter acidiphilus* was metabolically active at pH 2.9 and growth was observed after it increased to pH 3.5. ** For *Desulfosporosinus metallidurans* the complete oxidation pathway was suggested from its genome.

| Name | pH range optimum | °C range  optimum | Organic acid oxidation | Origin | Reference |
| --- | --- | --- | --- | --- | --- |
| *Thermodesulfobium*  *narugense^T^* | 4.0 – 6.5  ND | 37 – 65  50-55 | Incomplete | ARD  (sediment) | ^1^ |
| *Desulfosporosinus*  *acidiphilus^T^* | 3.6 – 5.5  5.2 | 25 – 40  30 | Incomplete | AMD  (sediment) | ^2^ |
| *Desulfosporosinus*  *acididurans^T^* | 3.8 – 7.0  5.5 | 15 – 40  30 | Incomplete | ARD (sediment) | ^3^ |
| *Thermodesulfobium*  *acidiphilum^T^* | 3.7 – 6.5  4.8 – 5.0 | 37-65 55 | Incomplete | Geothermal heated soil | ^4,5^ |
| *Desulfothermobacter*  *acidiphilus^T^* | 3.5* – 6.5  4.5 | 42-70 55 | Incomplete | Terrestrial hot spring  (sediment) | ^6^ |
| *Desulfosporosinus*  *metallidurans^T^* | 4.0 – 7.0  5.5 | 4-37 28 | Complete  (TCA)** | AMD (microbial mat) | ^7^ |
| *Acididesulfobacillus*  *acetoxydans^T^* | 3.8 – 6.5  5.0 | 25-42  30 | Complete (WLJ) | ARD  (sediment) | ^8^ |
