## Supplementary material for "Proton stress adaptation in acidophilic sulfate-reducing bacteria: insights from *Acididesulfobacillus acetoxydans* for acid mine drainage bioremediation": Table S17

**Table S17.** Overview of high-rate sulfate-reducing sulfidogenic bioreactors operated under acidic conditions. A,B,C Indicate that these rates were achieved within the same reactor run. Table adapted from ^1,2^. Acronyms: VSPR, volumetric sulfide production rate; SRB, sulfate-reducing bacteria; WWTP, waste water treatment plant; CSTR, continuous stirred-tank reactor; FBR, fluidized bed bioreactor; GLR, Gas-lift bioreactor; MBR, Membrane bioreactor. For the pH 3.5 VSPR all reactors are used, for pH 3.2 and 2.9 reactor R1 and R2 are used.

| **Reactor** | **pH** | **°C** | **Energy / Carbon source** | **Inoculum** | **VSPR [mg L^-1^ day^-1^]** | **Reference** |
| --- | --- | --- | --- | --- | --- | --- |
| GLR | 7.0 | 30 | H_2_/CO_2_ | Mixed SRB community | 10632 | ^3^ |
| FBR | 6.4 | 25 | Glucose + Acetate | WWTP sludge | 720^A^ | ^4^ |
| FBR | 4.5 | 30 | Glycerol | Acidophilic consortium | 710^B^ | ^5^ |
| MBR | 4.0 | 30 | Formate | Acclimated granular sludge | 5149 | ^6^ |
| FBR | 4.0 | 35 | Glycerol | Acidophilic consortium | 497 | ^7^ |
| MBR | 4.0 | 30 | H_2_/CO_2_ | Acclimated granular sludge | 205 | ^6^ |
| **CSTR** | **3.5** | **30** | **Glycerol** | ***A. acetoxydans*** | **102^C^** | This study |
| FBR | 3.2 | 25 | Glucose + Acetate | Acclimated WWTP sludge | 1992^A^ | ^4^ |
| **CSTR** | **3.2** | **30** | **Glycerol** | ***A. acetoxydans*** | **98^C^** | This study |
| FBR | 3.0 | 30 | Glycerol | Acidophilic consortium | 43^B^ | ^5^ |
| **CSTR** | **2.9** | **30** | **Glycerol** | ***A. acetoxydans*** | **63^C^** | This study |
